# Biological Process Activity Transformation of Single Cell Gene Expression for Cross-Species Alignment

**DOI:** 10.1101/555268

**Authors:** Hongxu Ding, Andrew Blair, Ying Yang, Joshua M. Stuart

## Abstract

The maintenance and transition of cellular states are controlled by biological processes. Here we present a gene set-based transformation of single cell RNA-Seq data into biological process activities that provides a robust description of cellular states. Moreover, as these activities represent species-independent descriptors, they facilitate the alignment of single cell states across different organisms.

---

The advent of single cell RNA sequencing (scRNA-Seq) technologies has greatly advanced our understanding of cellular states [1]. However, the signal-to-noise ratio of scRNA-Seq data is usually poor, confounding cellular state interpretation. Considering cellular states are controlled by genetic regulatory mechanisms [2], we propose using biological process activities (BPAs) in place of the expression of individual genes in the scRNA-Seq data, which leverages an ensemble of dozens of related genes. In this way, discrepancies in individual genes can be averaged out, yielding reproducible measurements unaffected by common technical noises such as batch effects [3] and drop-out events [4] (Figure 1B-D).

**Figure 1.**
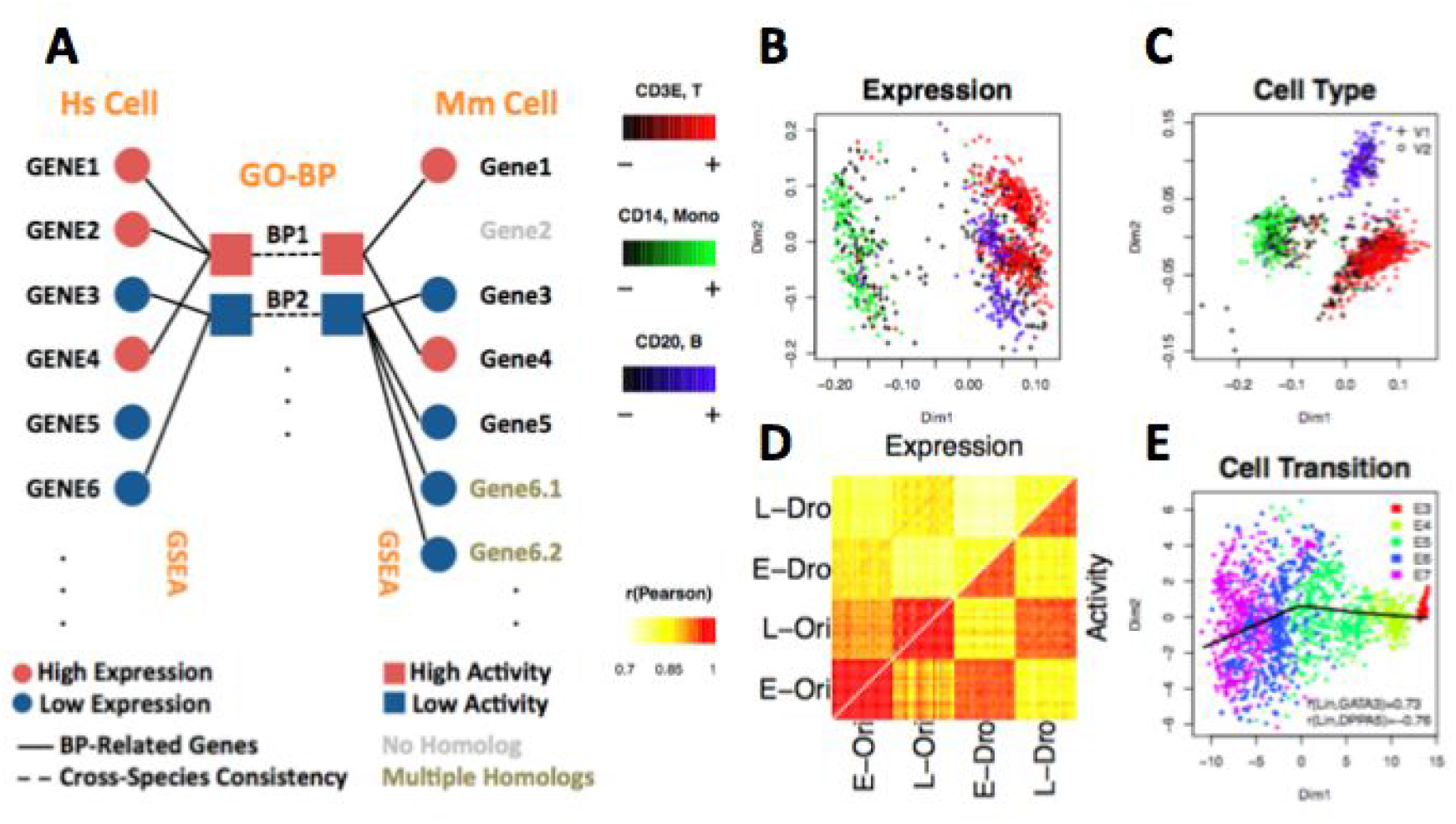
Biological process activity (BPA) transform overview and performance on single cell datasets. **(A)** Overview of biological process activity inference. Single cell gene expression profiles for human (outer left column) can be compared to a mouse gene expression profile (outer right column) using transformed biological process activity profiles for human (inner left) and mouse (inner right) even though the gene members of each Gene Ontology Biological Process (GO-BP) are distinct in each species (outer links). **(B)** Single peripheral blood mononuclear cells (PBMCs) profiled using 10x Genomics V1 and V2 chemistry were visualized using transcript expression features. Cells were color-coded according to expression of B-cell, monocyte and T-cell specific markers CD3E, CD14 and CD20, respectively. Two clusters for each cell type, corresponding to each of the Chromium chemistries are visible before the BPA transform. **(C)** Same as part B but PBMC data plotted after applying the BPA transform resulting in one cluster per cell type and no visible chemistry batch effect. **(D)** Drop-out events (Dro) were simulated into GTEx lung (L) and esophagus (E) bulk RNA sequencing (Ori) data (Supplementary Figure 2, see Methods); pairwise correlations between samples was computed and plotted resulting from comparisons using the original gene expression features (upper matrix triangle) and using BPA features (lower matrix triangle); high correlations, red; low correlations, yellow. BPA preserves same tissue comparisons even for drop-outs and has lower on average cross-tissue correlation than using gene expression features. (E) BPA transform preserves the developmental order of embryo development of the original study [13] based on the pseudotime inferred from principal curve [14] construction in t-SNE space [15] (Supplementary Figure 4).

Gene sets have been used extensively over the past few years to infer the activity of biological processes in many applications. The various catalogs of gene sets, e.g. Molecular Signature Database (MSigDB), group genes into categories of related function. Such gene sets allow particular pathways to be associated with the results of high-throughput assays. Gene set enrichment analysis (GSEA) is a particularly successful approach that summarizes the putative importance of a biological process using an ensemble of expression levels for a set of genes documented to play a role in a specific process [5].

We propose transforming gene expression levels into interpreted features using BPAs. The BPA transform is based on the published aREA method [6][7] that extends GSEA to interpret scRNA-Seq data as inferred process “activities”. Compared to previous studies [8][9], which used selected gene sets describing specific biological processes, e.g. cell cycle or TP53 pathway, for this study we use a comprehensive collection of gene sets, e.g. Gene Ontology (GO) Biological Process (BP), and the immunologic portion of the MSigDB collection called C7 [10], contributing ~650 and ~1800 gene sets after filtering, respectively (see Methods). BPA’s only option is to decide what gene sets to use for the transformation (e.g. gene sets can be selected based on their standard deviation, see Methods). For an individual cell and a specific pathway, the BPA transform runs an aREA-based single cell pathway enrichment analysis to create an activity score from the expression levels of the gene set members of the pathway (Figure 1A). In this way, the gene expression signature of an individual cell is converted to a BPA profile in which every pathway has a distinct activity score. Moreover, assuming the GO-BP terminology is consistent across species, gene set enrichment analysis can be used in each species separately to infer an activity for the same set of processes. Thus, inferences of activity for each category from single cell RNA-Seq data can be compared across species even though the categories between species have different gene members. In this way, datasets of human and model organisms can be combined directly to reveal functionally analogous cell types across species without the need to predict orthologs. We demonstrate the utility of using BPAs to align human and mouse datasets to shed light on their comparative and species-specific biology in early embryo development and in the cell types comprising the immune system.

Distinct, batch-specific clusters can be observed among peripheral blood mononuclear cell (PBMC) scRNA-Seq datasets when gene expression profiles are used directly (Figure 1B), but are no longer apparent after a BPA transformation (Figure 1C), giving comparable, or even better performance compared to batch-correction algorithms [3][11][12] (Supplementary Figure 1). In this example, the clustering of the PBMCs recapitulates the groups of B-cells, T-cells, and monocytes as denoted by the cell type-specific markers CD3E, CD14 and CD20, respectively. In addition, biological process activity is insensitive to drop-out events, illustrated through a controlled simulation in which drop-outs are introduced to mimic their distribution in real scRNA-Seq data. Complete RNA sequencing datasets of bulk tissue samples were taken from the GTEx lung (L) and esophagus (E) collection and labeled as “original” (Ori) while their counterparts containing simulated drop-out events were labelled “drop-out” (Dro) (Supplementary Figure 2, see Methods). We found that drop-out events appreciably decrease correlations within the same biological state. Moreover, correlations between different tissues with full data, e.g. r(L-Ori, E-Ori) was found to be higher than correlations between the same tissue type having drop-out data e.g. r(L-Dro, L-Dro) or r(E-Dro, E-Dro). Thus, artifacts in downstream analyses could be introduced when using transcript-level data containing drop-out events, since single cells will cluster according to the extent of the drop-out effect rather than the measured biological conditions. The inferred biological activity preserves within-tissue correlations, and reduces cross-tissue correlations (Figure 1C). Taken together, inferred BPA profiles produced clusters with distinctly enriched PBMC cell types according to marker gene expression (Figure 1D and Supplementary Figure 3), as well as the known ordering of state transitions in a human preimplantation embryo dataset (Figure 1E, Figure 2A and Supplementary Figure 4; see Methods).

**Figure 2.**
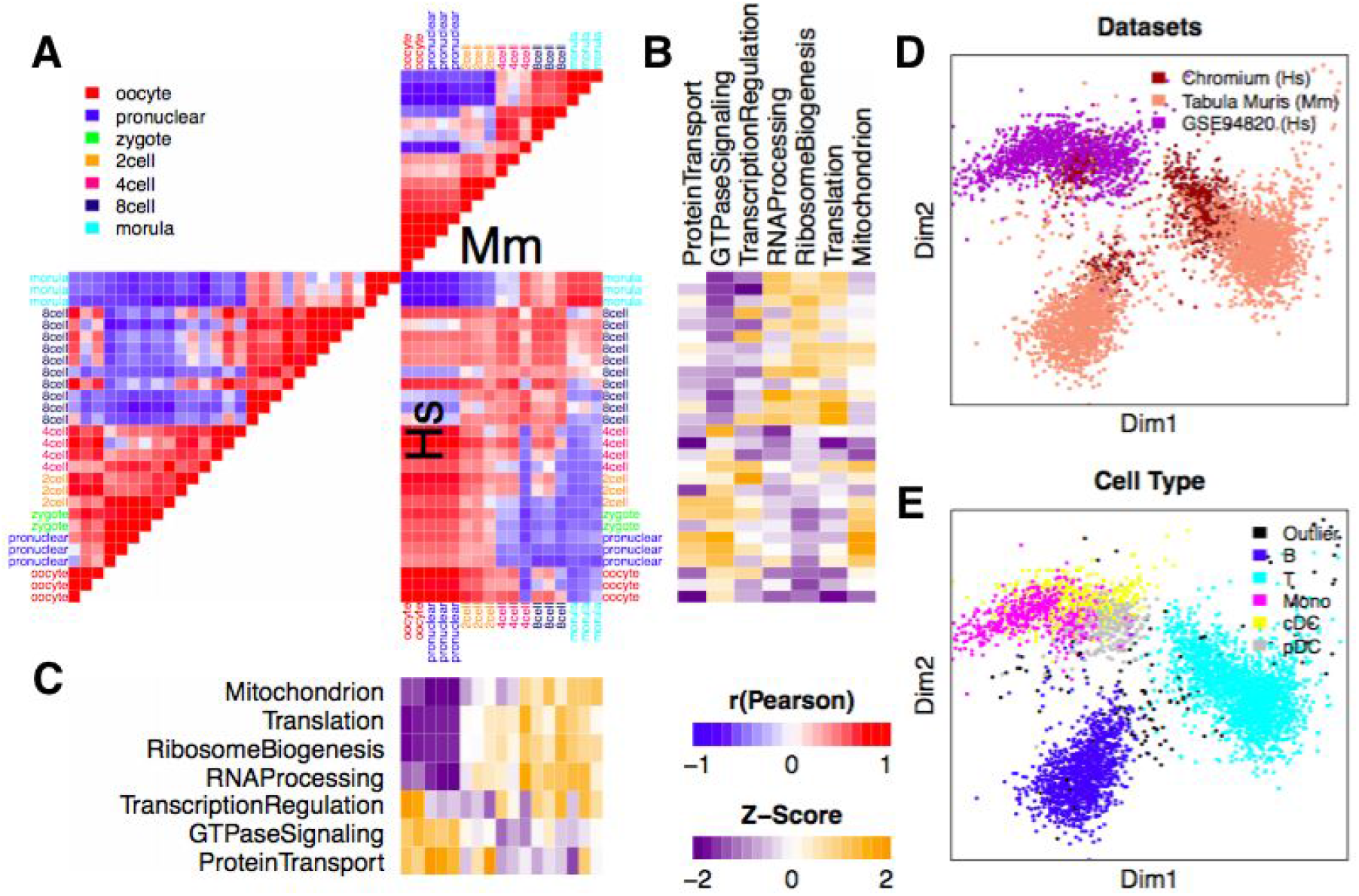
Aligning human and mouse single cell datasets using biological process activity (BPA). (A) Human and mouse early embryo single cells were taken from [2]. Pairwise correlation of cells shown as a heatmap; high, red; low, blue. (B) Inferred activity of biological processes described in the original study for the mouse dataset; high activities, yellow; low activities, purple. (C) Same as in part B but for the human dataset. (D) Multi-dimensional scaling (MDS) view of the cross-species BPA-integrated immune cell studies from human and mouse including two human studies -- Chromium, brown; and GSE94820, purple -- and one mouse study -- Tabula Muris, peach. (E) Same data as in part D showing the MDS view of the distinct immune cell types (colors) showing how cell types (e.g. T-cells, light blue) from both species cluster near one another.

We next performed cross-species single cell state alignment using BPA profiles in place of gene expression profiles. A previous effort used the expression pattern of one-to-one orthologous genes to align single cells across species [16]. However, as orthologs are usually determined by computational analysis of protein sequence alone [17], the expression pattern of orthologous genes may not be the same across species [18]. The BPA transform, on the other hand, provides a common set of terms from which detailed gene sets can be retrieved in a species-specific manner [10]. Each species can be analyzed separately using their species-specific gene sets and then merged across species at the BPA level assuming the pathway categories are equivalent, giving an overarching perspective of cellular states and transitions across various organisms.

We analyzed scRNA-Seq profiles reported in a human-mouse comparative study on embryo development. While early embryo development is a continuous process, it can be roughly divided into three steps [2]. By analyzing human and mouse single cells separately, we find that the three steps were recapitulated. In human, the first step spans the oocyte to 4-cell stage; the second step includes the 8-cell stage; and the third step includes only the morula stage. In mouse, the first step of the developmental timeline is relatively short compared to human, including the oocyte and pronuclear stages; the second step includes the 2-cell and 4-cell stages; and the last step includes the 8-cell and morula stages (Figure 2A). The data sets transformed with GO-BP produce the expected alignment of the three steps between the two species (Figure 2A), which was further confirmed objectively using dynamic time warping (Supplementary Figure 5) [16][19]. In addition, the stage-specific activation pattern of biological processes determined in the original study were recapitulated in both human and mouse cells (Figure 2B, C).

We next performed BPA analysis to compare and align human and mouse immune cells. We included scRNA-Seq profiles of human PBMCs from a healthy donor (Chromium, Figure 1A, C and Supplementary Figure 3), mouse spleen and thymus (Tabula Muris, Supplementary Figure 3) [20] and human monocytes and dendritic cells (GSE94820, Supplementary Figure 3) [21]. To extend the scope of biological process selection, and to better describe the cellular states, we also included an immunologic gene set (see Methods). Within each individual dataset, cell types (Supplementary Figure 3), as well as cell type-specific biological processes (Supplementary Table 1-3) were recapitulated using BPA, benchmarking the immunologic gene set in interpreting cellular states. Data sources, as well as cell types, for the integrated analysis of the three datasets are shown in Figure 2D-E. Although some species-specificity was observed, cells are primarily clustered according to cell types, forming clustered populations of T-cells and phagocytes (composed of monocytes and dendritic cells) contributed by both species.

To test the significance of this result, we used an analysis of variance to measure the separation of cells of the same cell type to those of the same species in the transformed data. Specifically, we calculated the variance within cell types (Vt) and the variance within species (Vs) and found that the ratio (Vt/Vs) was indeed significantly higher than unity (P=5.0×10^−12^, F-test, See Methods). In contrast, mapping genes to orthologs and using orthologous gene expression to combine the datasets resulted in a significantly lower level of species mixing within cell types (F-test P=0.0035). Visual inspection of the hierarchical clustering dendrogram of the BPA result (Supplementary Figure 6) shows that the major division of the cells falls along the axis defining the B, T, and monocyte cell types and the human-mouse divisions occur as minor splits further down the tree. In contrast, clustering the ortholog gene expression data produces a dendrogram with a major split that falls along the species and dataset distinctions. These results indicate that single cells from different experiments and across species can be combined effectively using BPA.

To evaluate the performance of BPA analysis in separating closely- and distantly-related species, we integrated human (E-MTAB-3929) [13], zebrafish (GSE66688) [22] and mouse (GSE65565) [23] embryo-related single cells. Single cells from human preimplantation embryos, especially in the early stages (E3 and E4), overlapped with mouse embryonic stem cells (Supplementary Figure 7). This finding is consistent with the mouse ESCs being derived from blastocyst, an early-stage pre-implantation embryo. Single cells from human preimplantation embryos and mouse ESCs are separated from single cells of the zebrafish embryo, indicating zebrafish are less related to the cells of human and mouse. On the other hand, due to different scRNA-Seq platforms, gene expression profiles of the three datasets are not comparable. Therefore expression-based Seurat [24] analysis failed in cross-species integration (Supplementary Figure 8). This further confirms that the BPA transform is resilient to common “technical batch effects” among single cell expression profiles.

In summary, we have presented a gene set enrichment analysis-based approach for inferring biological process activities (BPAs) within single cells. Transforming the gene-level data into interpreted features representing cellular processes produces a dataset that is less influenced by common technical noises in scRNA-Seq profiles. The transformed data preserves the integrity of cellular states and their transitions. Moreover, analysis in BPA space enables a straightforward comparison of cell states across platforms and species with very few parameter settings and without the need for predicting orthologs. Using the BPA approach, model organism datasets can be directly combined with a human counterpart to uncover inter-species commonalities and differences in evolution, normal development, and diseases at the resolution of individual cells. Other transformations of the data are possible that could lead to similar benefits. For example, a master regulator analysis approach like VIPER [6][7], which infers the activity of transcription factors from their targets, might also be used. Other transforms such as deep autoencoders [27] or network diffusion [28] are also possible. Methods that preserve the explicit association with genes (e.g. VIPER and network diffusion) can be run prior to BPA transformation and thus offer the potential for exploring a combination of approaches. Finally, in addition to single cell datasets, BPA is applicable to bulk expression analysis (Supplementary Figure 9), serving as a general approach for describing and combining biological states across datasets and species.

## METHODS

### Gene set selection and filtering

Gene sets were downloaded from the CRAN R msigdbr package that includes the MSigDB (http://software.broadinstitute.org/gsea/msigdb/index.jsp) dataset [5]. In this study, the biological process subset of MSigDB’s GO gene sets (C5) and the MSigDB immunologic gene sets (C7) for *Homo sapiens, Mus musculus* and *Danio rerio* were used in the presented analyses. C7 contains paired up-regulated and down-regulated gene sets. Since single-cell RNA-sequencing profiles have high drop-out rates, the accuracy for quantifying under-expressed genes is low. To avoid using such genes, as well as redundancy caused by paired gene sets, we retrained only the up-regulated gene sets in C7 for analyses in the study. The gene sets were filtered by the number of genes in the gene set since aREA-based BPA tends to assign higher activities to larger gene sets. On the other hand, smaller gene sets yield less robust activities as indicated by an increase in the coefficient of variance as gene set sizes decrease (Supplementary Figure 10). For the C5-based gene sets, we mitigated these biases by restricting the size of the gene sets to range between 50 and 100 genes. For the C7 gene sets, we controlled for size by selecting gene sets ranging between 190 and 210. After filtering, for C5, 685, 680 and 722 gene sets were kept in mouse, human and zebrafish, respectively. For C7, 1741 and 2156 gene sets were kept in mouse and human, respectively (Supplementary Figure 11).

### Simulating drop-out effects in RNA-Seq data

We randomly selected 20 samples from each of the GTEx lung and esophagus bulk RNA sequencing samples. To mimic the single-cell RNA sequencing scenario, the simulated drop-out rate was determined by using a drop out probability that is a function of the expression level of each transcript. For example, more lowly expressed transcripts have a higher likelihood of drop-out than those that are more highly expressed. Such a relationship was determined empirically by analyzing Chromium and Ginis *et al*. [18] datasets. Although the drop-out rate varies depending on the cell type, such a relationship still holds. The simulation was done in lung and esophagus datasets separately, yielding 81.32% and 82.84% overall drop-out rate (Supplementary Figure 2).

### Biological Process Activity (BPA) Transformation of single cell RNA-Seq Data

Single sample GSEA (ssGSEA) was used to quantify activity profiles from the original gene expression [25]. ssGSEA was performed using the aREA() function from the Bioconductor R viper package. The aREA() function performs a rank-based enrichment analysis, which provides a computationally efficient analytical approximation of the widely-used GSEA tool [6][7]. Since the normalized enrichment scores given by the aREA() function are essentially z-scores they can be directly merged across datasets without any further normalization.

The original gene expression is provided as the input signature to ssGSEA, to quantify a measure of “activity” for each biological process. This contrasts the typical use of single cell GSEA analysis that quantifies a relative activity by subtracting the average expression signature of the entire dataset from the expression of each individual cell [6][7]. In this study, we used the log-transformed original single cell expression as the signatures for inference instead of using a differential measure that would strongly depend on the composition of the dataset and possibly impact cross-dataset integration (Supplementary Figure 12).

Many genes occur in multiple pathways, creating redundancies among the gene sets due to the overlap of their members. However, we found that down weighting gene sets according to their overlaps using “shadow analysis” in the Bioconductor R viper package [6][7] produced BPAs with very little difference from their unweighted versions (see Supplementary Figure 13). For this reason, we chose to keep all of the genes within a biological process to insure a comprehensive representation and to use the unweighted gene sets to analyze the data in this study.

We tested the sensitivity of BPA to select relevant pathways using a dataset with known cell types. Our assumption is that relevant pathways should contain some genes with expression levels that vary between cells of different cell types. Thus, the standard deviation (SD) across all cells may reflect that a gene set encodes cell type-related information. As shown in Supplementary Figure 14, within the heterogeneous Chromium 10x PBMC dataset, the gene sets in the MSigDB collection gave overall higher SD values compared to random gene sets (negative control), indicating that some pathway sets reflect immune cell-related information. This suggests that the SD values could be used to select specific biological processes to analyze a given dataset. In support of this, context-matching biological processes of the immune system (C7) gave higher SD values than general pathway gene sets (C5) on average for the PBMC dataset (Supplementary Figure 15).

### Significance of cell type separation, dimensionality reduction, and clustering

We used a statistical test to assess the degree to which datasets of different species are concordant either in the original gene expression space or upon transformation using BPA. Intuitively, cells of the same cell type but from different species should overlap thus maintaining the separation of distinct cell types yet mixing the species. To quantify this notion, we used an analysis of variance in which we measured the variance within species (*V_s_*) and within cell types (*V_t_*) and calculated their ratio *V_t_*/*V_s_*. Values higher than unity indicate cells of the same cell types are closer together (lower variance) compared to a cell of a different cell type from the same species (higher variance). The ratio follows an F-distribution with (*N-M_t_*) and (*N-M_s_*) degrees of freedom where *N* is the number of cells, *M_t_* is the number of cell types (*M_t_=3* for the PBMC dataset) and *M_s_* is the number of datasets (*M_s_*=3, including PBMC, Tabula Muris and GSE94820 for the human-mouse analysis). To combine the original gene expression vectors across species without transforming with BPA, we mapped all genes from human and mouse to their orthologous counterparts using the human-mouse mapping table available in the R Bioconductor biomaRt package.

To visualize both the BPA transformed and original gene expression spaces, we produced a multidimensional scaling (MDS) dimensionality reduction using the cmdscale() function in the R stats package. To detect cell types, DBSCAN [26] clustering on the 2D MDS space was performed using the dbscan() function in the R CRAN dbscan package. To be consistent with the pseudotime analysis of the original study [13], we used the 2D t-SNE [15] using the Rtsne() function from the R CRAN tsne package, followed by pseudo-lineage analysis using principal curves [14]. Principal curves were calculated using the principal.curve() function from the CRAN R princurve package. We also performed hierarchical cluster analysis on the datasets to produce the dendrograms illustrating the BPA data divides cell type more significantly than species compared to using the original gene expression data associated with predicted orthologs.

### Data availability

GTEx bulk RNA sequencing profiles can be found from the website: https://gtexportal.org/home/. We downloaded the provided normalized expression profiles and log-transformed them into log2(RPKM+1) for downstream analysis.

Mouse and human esophageal epithelium normalized bulk RNA-Seq expression profiles: https://www.ncbi.nlm.nih.gov/geo/query/acc.cgi?acc=GSE116272.

scRNA-Seq profiles for the human PBMC dataset were taken from healthy donors generated using 10x Genomics V2 and V1 chemistry and available from: https://support.10xgenomics.com/single-cell-gene-expression/datasets/2.0.1/pbmc4k, https://support.10xgenomics.com/single-cell-gene-expression/datasets/1.1.0/pbmc3k. We downloaded the provided UMI counts and normalized by the sequencing depth as log2(TPM+1) for downstream analysis.

scRNA-Seq profiles for the human preimplantation embryo dataset, including time point: https://www.ebi.ac.uk/arrayexpress/experiments/E-MTAB-3929/. We downloaded the provided normalized expression profiles and log-transformed them into log2(RPKM+1) for downstream analysis.

scRNA-Seq profiles for the mouse embryonic stem cell dataset: https://www.ncbi.nlm.nih.gov/geo/query/acc.cgi?acc=GSE65525. We downloaded the provided normalized expression profiles and log-transformed them into log2(TPM+1) for downstream analysis.

scRNA-Seq profiles for the zebrafish embryo dataset: https://www.ncbi.nlm.nih.gov/geo/query/acc.cgi?acc=GSE66688. We downloaded the provided normalized expression profiles and log-transformed them into log2(TPM+1) for downstream analysis.

scRNA-Seq profiles for the human and mouse early embryos, including time point annotations: https://www.nature.com/articles/nature12364. We downloaded the provided normalized expression profiles and log-transformed them into log2(RPKM+1) for downstream analysis.

scRNA-Seq profiles for the human monocytes and dendritic cells, including cell type annotation: http://science.sciencemag.org/content/356/6335/eaah4573. We downloaded the provided normalized expression profiles and log-transformed them into log2(TPM+1) for downstream analysis.

*Tabula Muris* datasets: https://www.nature.com/articles/s41586-018-0590-4. We downloaded the provided counts of spleen and thymus datasets and normalized by the sequencing depth as log2(CPM+1) for downstream analysis.

All relevant data and analysis results are available from the authors.

### Code availability

All scripts are available at https://github.com/hd2326/BiologicalProcessActivity. The code contains all 7 categories of MSigDB gene sets (C1-7), including positional gene sets (C1), curated gene sets (C2), motif gene sets (C3), computational gene sets (C4), GO gene sets (C5), oncogenic gene sets (C6) and immunologic gene sets (C7) from 11 species including *Bos taurus, Caenorhabditis elegans, Canis lupus familiaris, Danio rerio, Drosophila melanogaster, Gallus gallus, Homo sapiens, Mus musculus, Rattus norvegicus, Saccharomyces cerevisiae* and *Sus scrofa*.

## AUTHOR CONTRIBUTIONS

H.D. and J.M.S. conceived and initiated the project. H.D., A.B. and Y.Y. collected public datasets and performed the analysis. H.D., A.B. and J.M.S. prepared the manuscript.

## COMPETING INTERESTS

All authors declare no competing interests.

## Notes

https://github.com/hd2326/BiologicalProcessActivity

https://gtexportal.org/home/

https://support.10xgenomics.com/single-cell-gene-expression/datasets/2.0.1/pbmc4k

https://support.10xgenomics.com/single-cell-gene-expression/datasets/1.1.0/pbmc3k

https://www.ebi.ac.uk/arrayexpress/experiments/E-MTAB-3929/

https://www.nature.com/articles/nature12364

http://science.sciencemag.org/content/356/6335/eaah4573

https://www.nature.com/articles/s41586-018-0590-4

