## Supplemental Information for "Biological Process Activity Transformation of Single Cell Gene Expression for Cross-Species Alignment"

### Supplementary Information

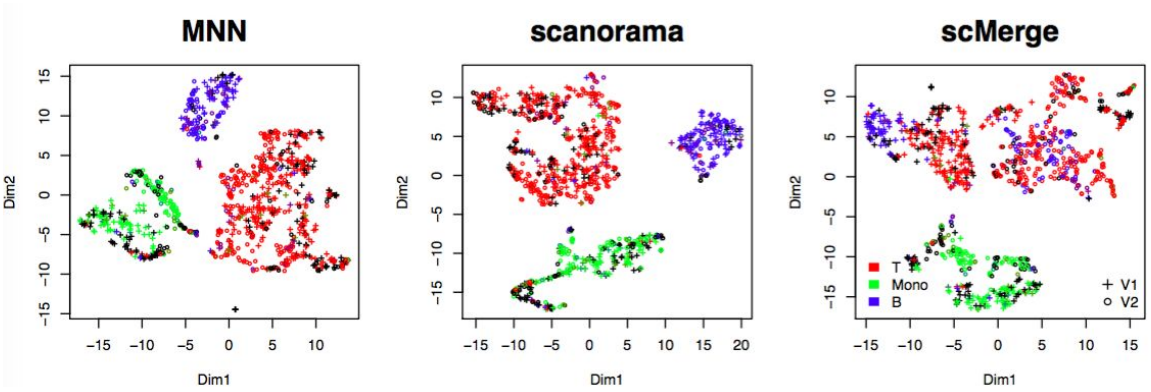


**Supplementary Figure 1.** MNN [Haghverdi2018], scMerge [Lin2018], and Scanorama [Hie2018] batch correction on the PBMC dataset. For the three algorithms, correction and visualization were performed using corresponding default parameters. As shown, Scanorama significantly reduced the “batch effect”, producing similar results as BPA analysis. Minor “batch effect” can be observed after MNN or scMerge transformation, with scMerge seeming to lose the resolution of separating B and T cells.

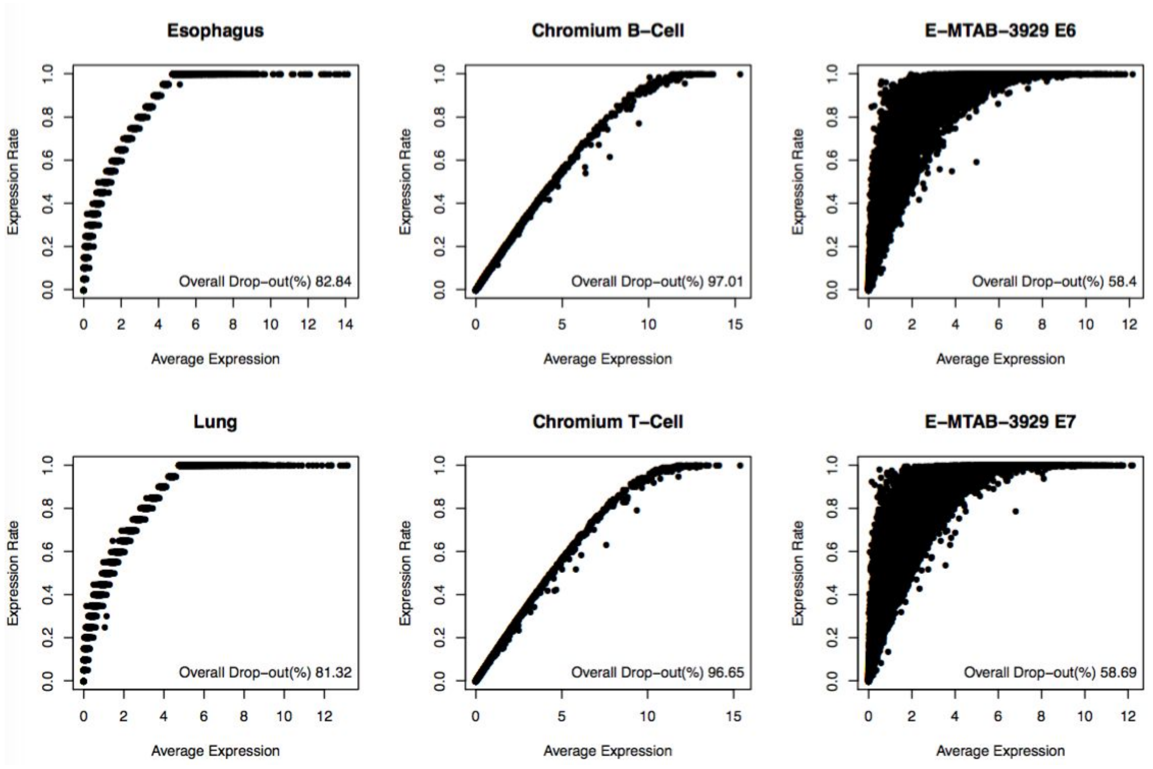


**Supplementary Figure 2.** Simulating drop-out effect from GTEx esophagus and lung bulk RNA sequencing profiles (see METHODS), as well as single cell drop-out effect from Chromium and [Petropoulos2016] datasets.

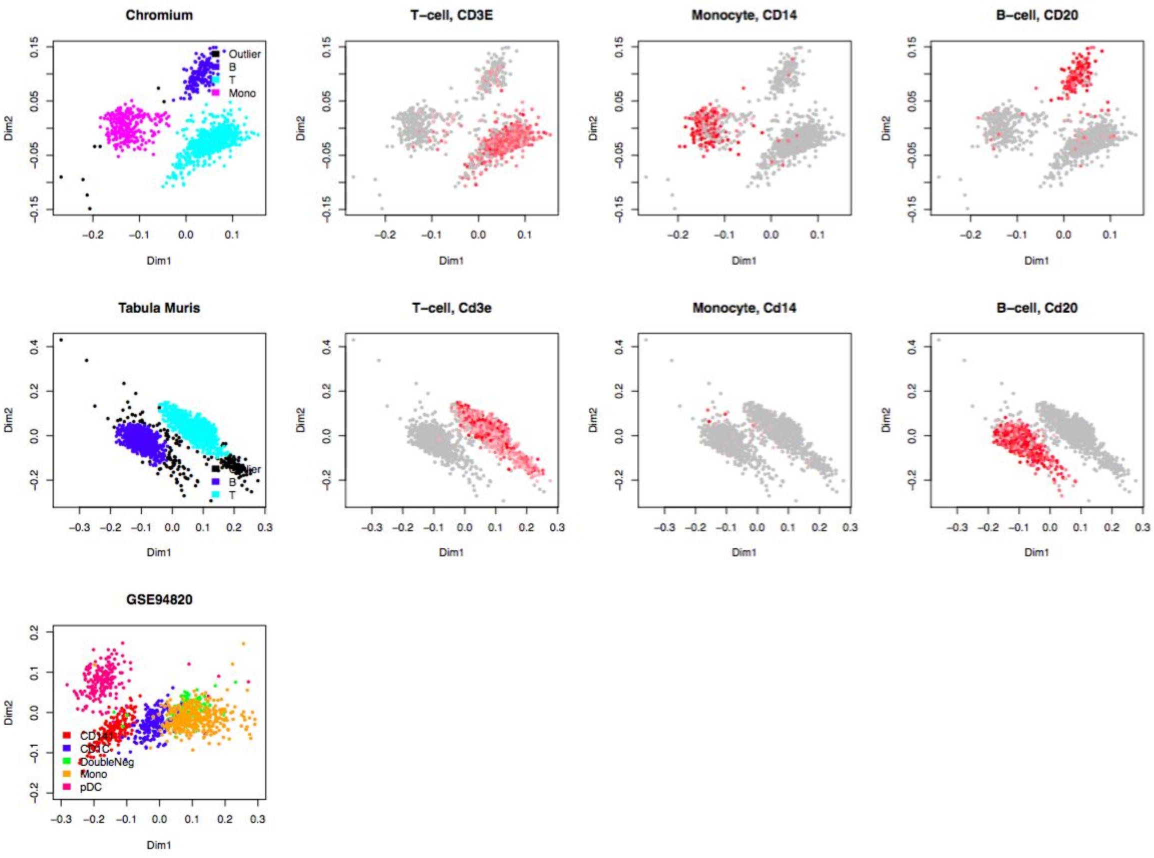


**Supplementary Figure 3.** Clustering analysis and marker expression annotation of single-cell RNA sequencing datasets (see METHODS). The annotation of dataset GSE94820 (human monocyte-dendritic cell single-cell RNA sequencing profiles) was provided by the original study [Villani2017].

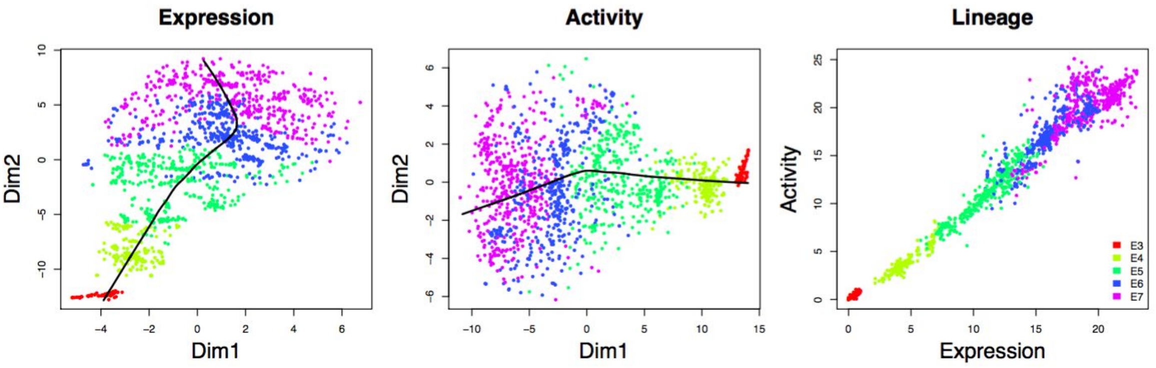


**Supplementary Figure 4.** Pseudo-lineage analysis on [Petropoulos2016] dataset at gene expression and biological process activity level (see METHODS). Expression- and activity-based pseudo-lineages have strong correlation, and both consistent with time point annotation.

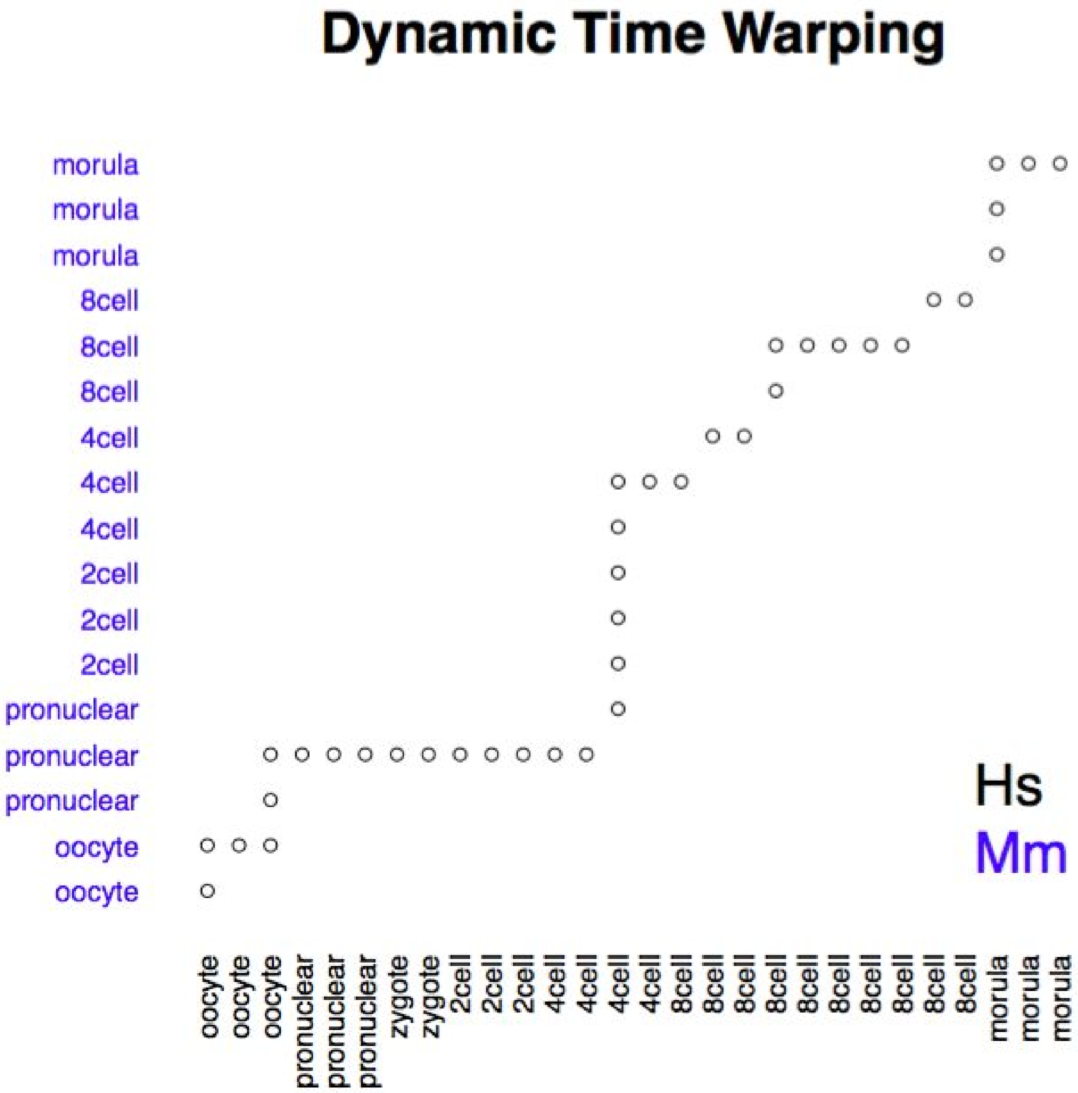


**Supplementary Figure 5.** Dynamic time warping-based cell aligning analysis [Alpert2018, Sakoe1978] between human and mouse early embryo single cells in [Xue2013] using pairwise correlation computed from biological process activity profiles. Dynamic time warping was performed using dtw() function in CRAN R dtw package.

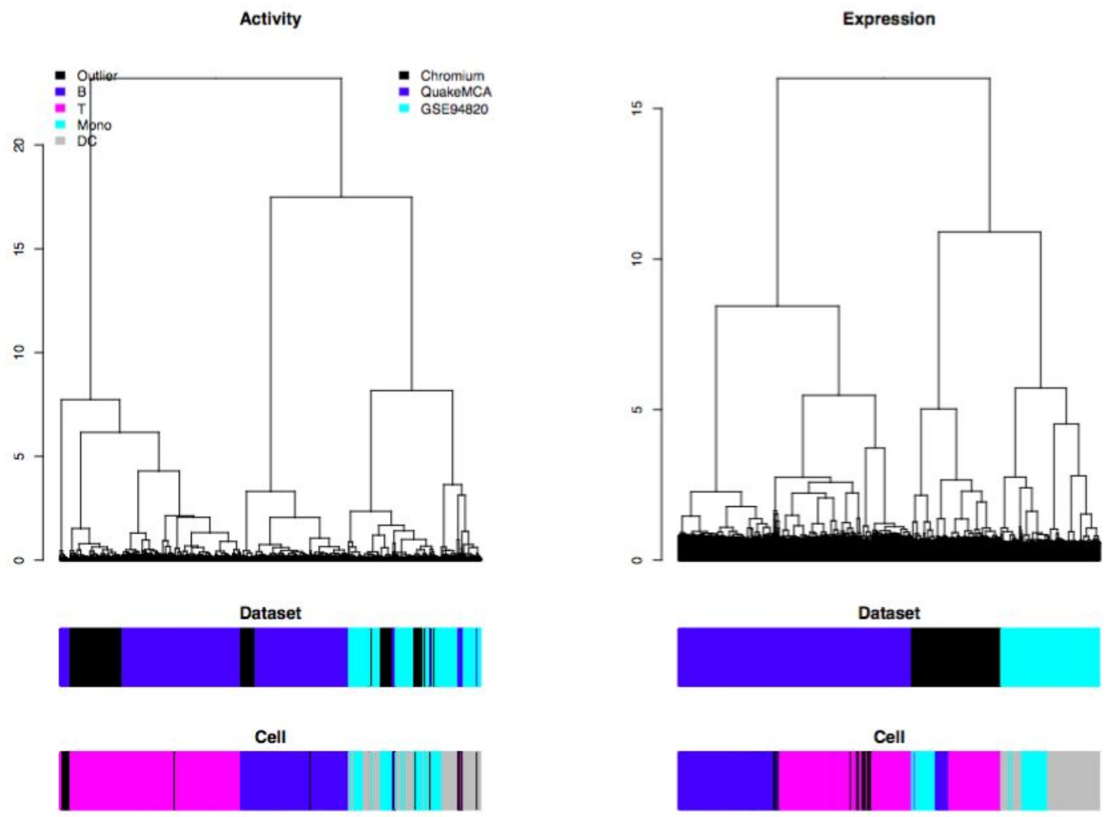


**Supplementary Figure 6.** Dendrogram representations of human-mouse cross-species immune cell analysis for the BPA transform (left dendrogram) and ortholog gene expression (right, see METHODS). For BPA, the major branch separates the data by the three major cell types -- B, T and monocyte-dendritic cell lineages (lower annotation bar with B-cells, blue; T-cells, pink; Monocytes, light blue; dendritic cells, gray; and DBSCAN-identified outliers, black). For ortholog expression, the major division falls along the species and dataset (top annotation bar in each plot, Chromium, Hs, black; Tabula Muris, Mm, blue; GSE94820, Hs, light blue).

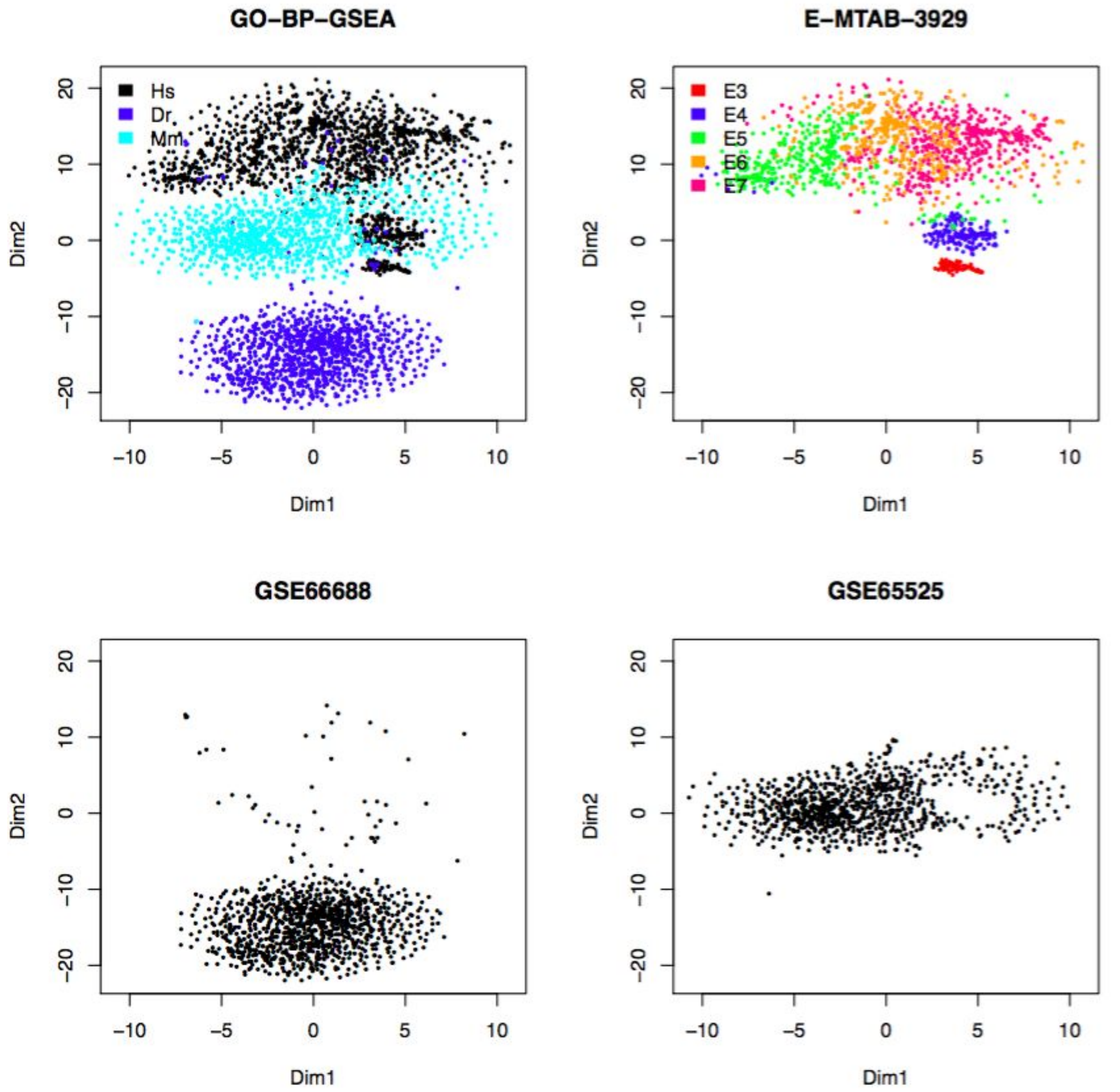


**Supplementary Figure 7.** Integrating human (Hs, E-MTAB-3929, E3-7 stage annotation provided by original study) [Petropoulos2016], zebrafish (Dr, GSE66688) [Satija2015] and mouse (Mm, GSE65525) [Klein2015] embryo-related cells, evaluating the performance of biological process activity analysis in separating closely- and distantly-related species.

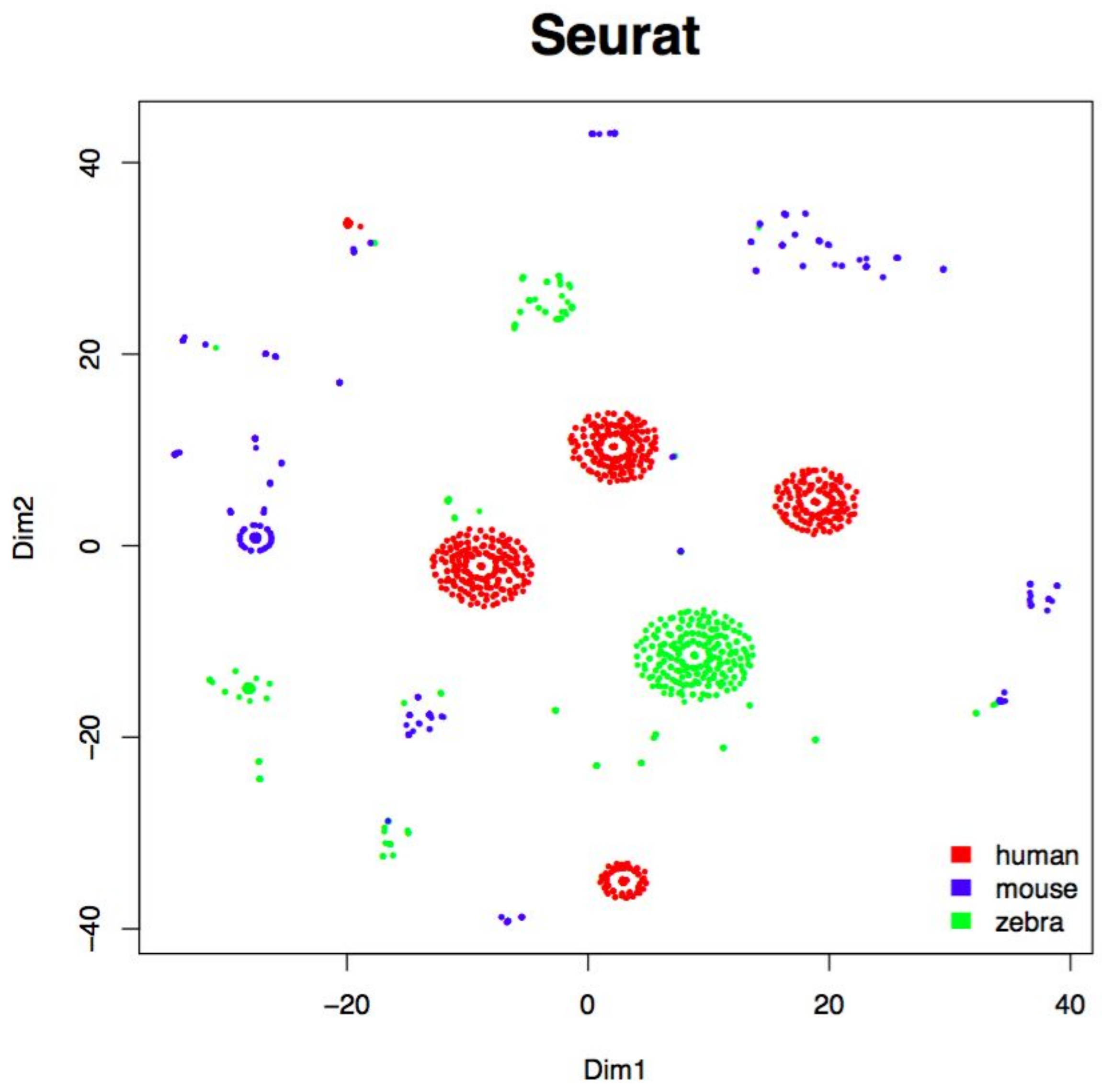


**Supplementary Figure 8.** Cross-species integration of datasets described in Supplementary Figure 7 using Seurat [Butler2018]. Mouse and zebrafish datasets were “humanized” by orthologous gene maps extracted using Bioconductor R biomaRt package. Cross-species integration and result visualization were derived with default Seurat parameters. Sequencing platforms: human, full-length scRNA-Seq without UMI; mouse, 3’ scRNA-Seq with 6nt UMI; zebrafish, 3’ scRNA-Seq with 5nt UMI. Expression quantification: human, RPKM; mouse, TPM; zebrafish: TPM.

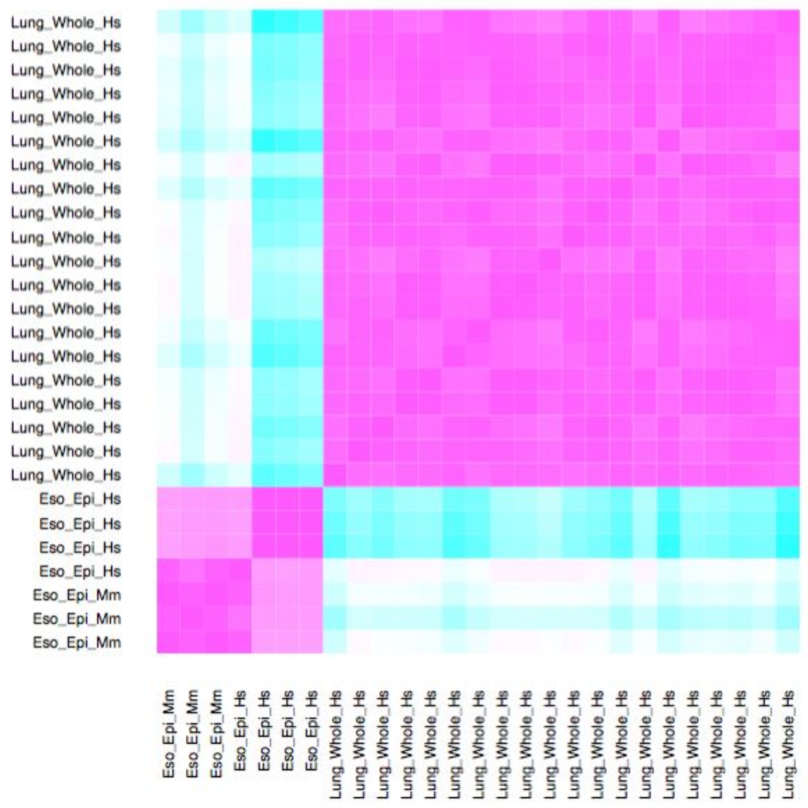


**Supplementary Figure 9.** Applying BPA analysis to bulk expression profiles, for analyzing biological states in human and mouse. Mouse esophageal epithelium (Eso_Epi_Mm) and human esophageal epithelium (Eso_Epi_Hs) profiles were obtained from [Zhang2018]. Human whole lung profiles were obtained from GTEx (see Figure 1) clustered separately from esophageal epithelium profiles, serving as “negative control”.

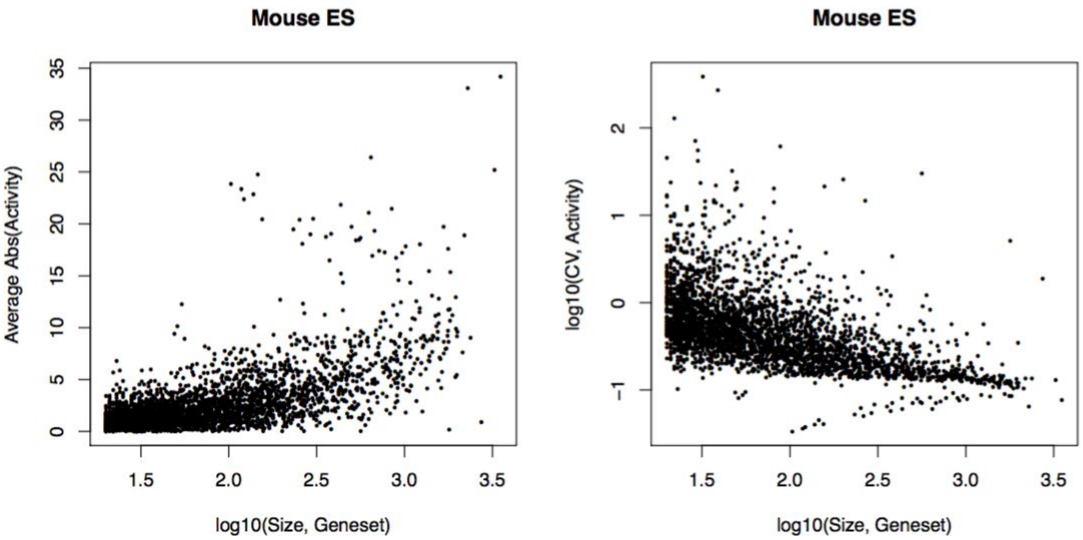


**Supplementary Figure 10.** Gene set size related bias in BPA analysis. Larger gene sets can give higher inferred activity, while inference from smaller gene sets is less robust, indicated by increased coefficient of variance (CV). The analysis was performed in a homogenous mouse embryonic stem (ES) cell line [Klein2015] to exclude the possibility that the variability is not caused by the presence of different cell types.

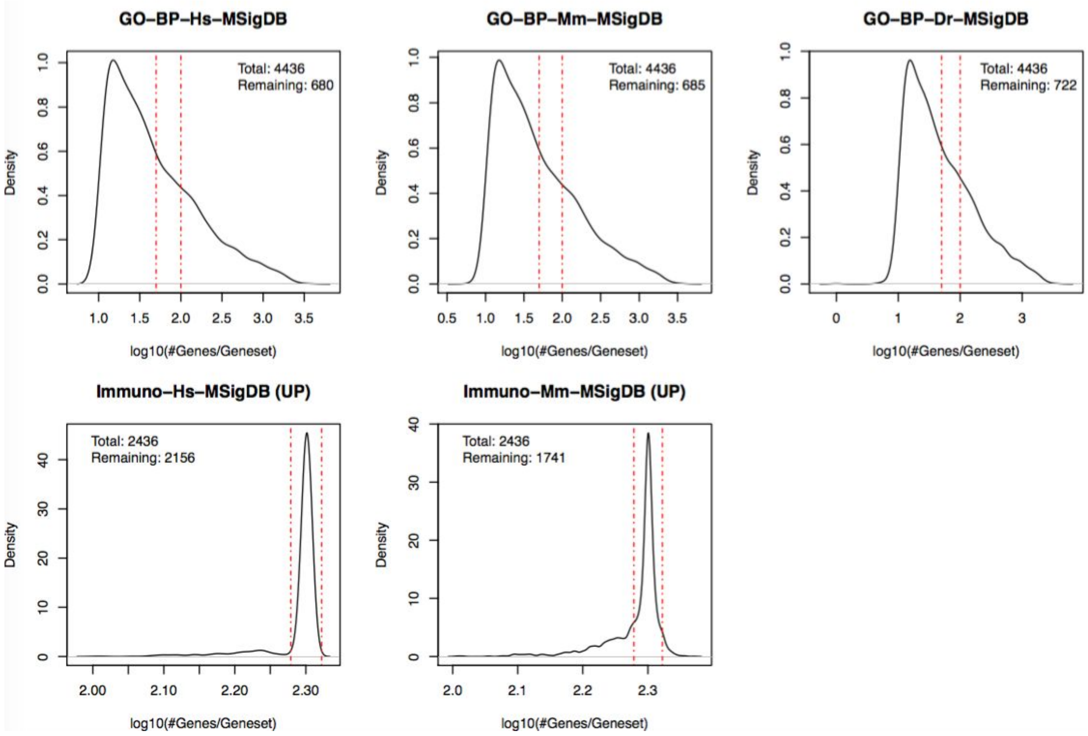


**Supplementary Figure 11.** Size-based gene set selection. Size probability density distribution of included gene sets, with the vertical red dashed lines showing the gene set size cut-off used for BPA analysis. For the immune gene sets, only annotated up-regulated (UP) gene sets were included.

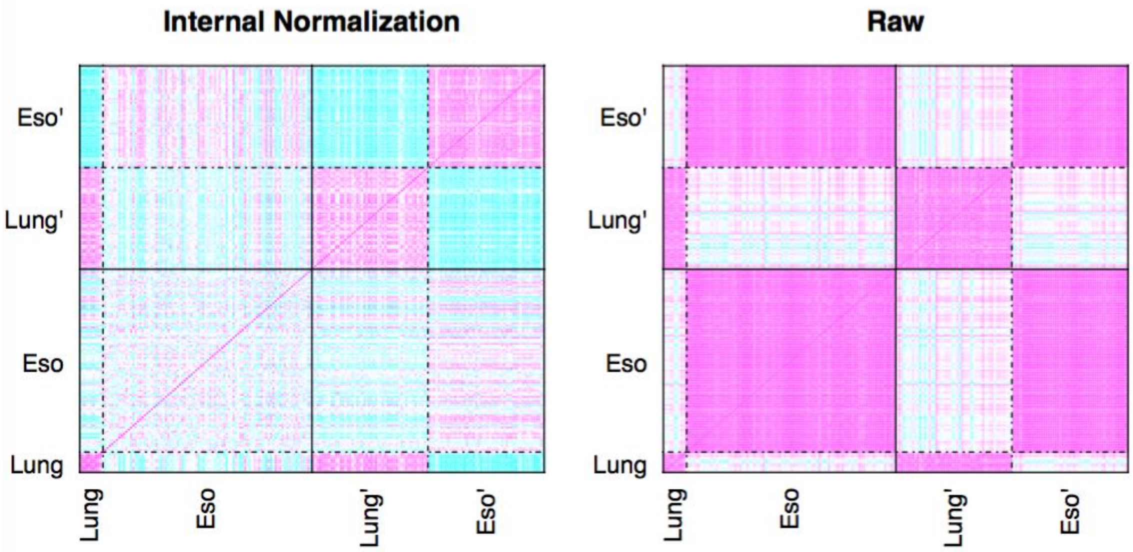


**Supplementary Figure 12.** Cross-dataset integration might be biased by different ways of generating expression signature. Symmetric sample-by-sample distance matrices in which an “internal normalization” was used to derive relative expression (left matrix) compared to using the original “raw” expression levels (right matrix). For “internal normalization” the “double-rank transformation” [Alvarez2016][Ding2018] was used. For this, the genes were ranked according to their expression within each individual cell, followed by extracting gene-wise median and median absolute deviation (mad). We then calculate (rank - median)/mad, as the single cell gene expression signature. In this case, each single cell will be internally normalized to the “average” of the entire dataset and provided to the BPA analysis producing a “relative activity”. For the “raw” result, absolute gene expression profiles quantified by log2(TPM+1) were passed to the BPA analysis, producing “absolute activity” vectors for each cell. From GTEx lung and esophagus datasets, we randomly selected i) 10 lung + 90 esophagus (Lung, Eso), and ii) 50 lung + 50 esophagus (Lung’, Eso’) samples to perform BPA analysis. For “internal normalization”, correlation within Eso samples within Lung+Eso dataset is much lower on average. This is because the dataset is dominated by Eso, therefore internal normalization emphasizes differences between Eso samples. While for Lung’+Eso’ dataset, which has a more balanced sample number between the two tissue types, internal normalization captures differences between Lung’ and Eso’, In this case, when doing cross-dataset integration between Lung+Eso and Lung’+Eso’, due to inter-sample correlation, Eso and Eso’ will not be combined, causing artifact.

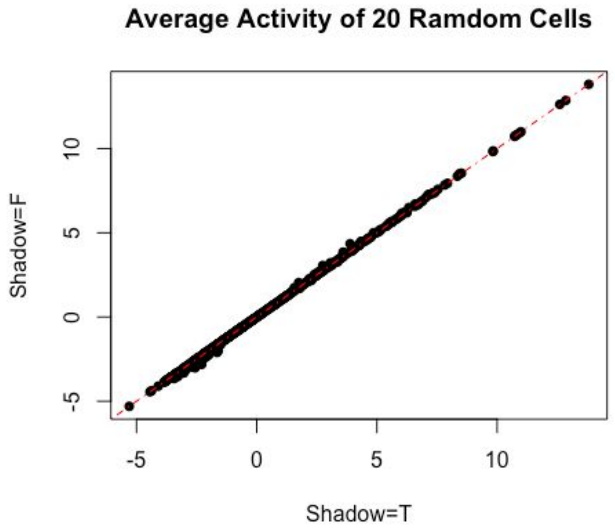


**Supplementary Figure 13.** Overlapping gene sets have little influence on BPA analysis. “Shadow analysis” was used to mitigate the influence of overlapping gene sets using the Bioconductor R viper package. Analysis was performed in 20 randomly selected cells from [Klein2015] homogenous mouse embryonic stem (ES) dataset both with (X-axis) and without (Y-axis) the shadow analysis illustrating the derived activities are highly similar from these two versions of the analysis.

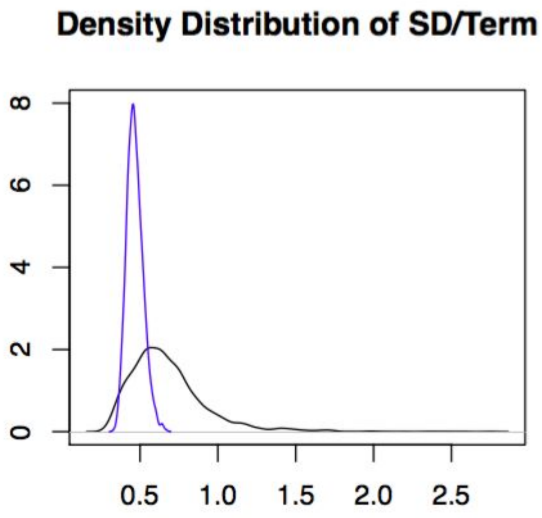


**Supplementary Figure 14.** The activities of pathways have higher standard deviations (SD) across the Chromium 10X PBMC dataset compared to random gene sets, reflecting the sensitivity of BPA analysis. BPAs were inferred for each pathway in from the GO-BP (C5) and Immune-related (C7) collections and their standard deviations were calculated and plotted, black curve. Similarly, the BPAs for random gene sets, created to match the sizes of the original pathways, were calculated and their standard deviations across the dataset plotted, blue curve.

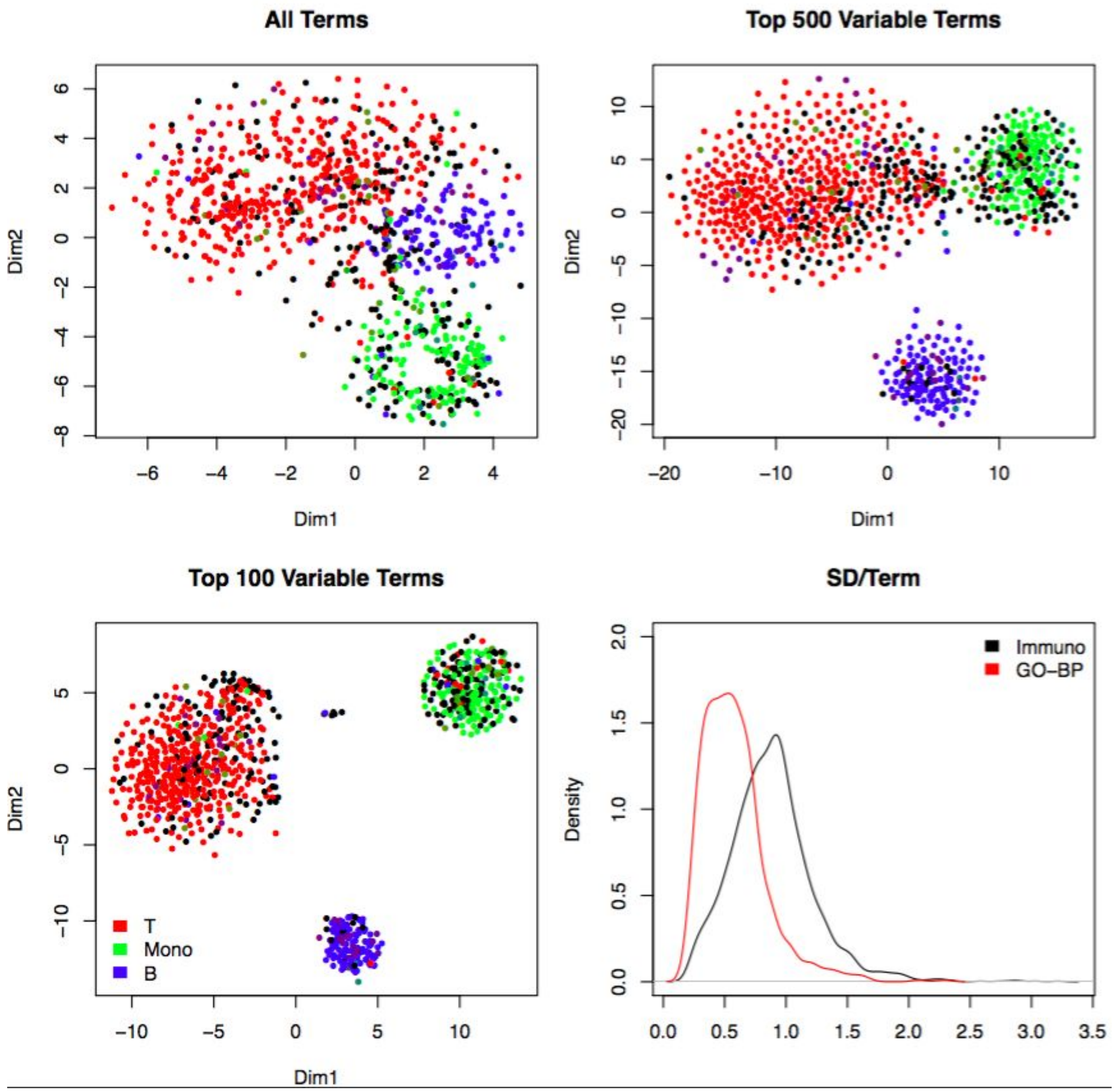


**Supplementary Figure 15.** Prioritizing informative biological processes using standard deviation (SD) values. Activity profiles were inferred from heterogenous Chromium 10x PBMC dataset, with combined C5 and C7 sets. Using biological process gene sets with high SDs gave clear separation of cell populations. Upper-left: BPA transformed datasets using all gene sets from C5 and C7. Upper-right: BPA transformed data using top 500 gene sets as scored by their BPA standard deviations. Lower-left: BPA transformed data using top 100 gene sets as scored by their BPA standard deviations. Lower-right: The immune-related (C7) gene sets gave higher SD values on average than other gene sets (in GO-BP), indicating that informative and relevant gene sets provide informative information upon BPA transformation.

**Supplementary Table 1.** Sorted combined GO biological process (C5) and immunologic (C7) gene set names according to differential activity in B-cells, T-cells and monocytes in Chromium 10x dataset.

**Supplementary Table 2.** Sorted combined GO biological process (C5) and immunologic (C7) gene set names according to differential activity in B-cells and T-cells in [TabulaMurisConsortium2018] dataset.

**Supplementary Table 3.** Sorted combined GO biological process (C5) and immunologic (C7) gene set names according to differential activity in monocytes, pDCs and cDCs in [Villani2017] dataset.
